# Label-free Raman imaging defines distinct cell populations in human skin

**DOI:** 10.64898/2026.02.18.706712

**Authors:** Kaori Sugiyama, Masahiro Ando, Mizuho Ishikawa, Aiko Sada, Haruko Takeyama

## Abstract

Understanding cellular heterogeneity in human skin is crucial for regenerative medicine and tissue engineering. In this study, we applied label-free Raman imaging to visualize molecular features corresponding to the three-dimensional architecture of the epidermis. Spatially resolved Raman spectra, combined with multivariate data analysis, enabled the identification of cell-layer–specific molecular signatures. Based on the region-specific spectra analysis, component C5 was predominantly localized to the basal layer within rete ridges and was characterized by β-sheet–enriched keratin features. This spatially restricted distribution reflects the molecular microenvironment of epidermal stem cell niches, suggesting that C5 may serve as a biomarker for basal stem cell populations associated with skin undulations. These findings provide insight into the molecular basis of epidermal architecture and demonstrate the potential of Raman spectroscopy as a label-free tool for evaluating stem cell localization and differentiation status.

## Introduction

Advances in biomedical imaging have deepened our understanding of tissue biology by enabling the *in situ* investigation of cellular and molecular processes. Raman spectroscopy is a powerful, noninvasive, label-free technique that can identify and spatially map molecular species via intrinsic vibrational spectra. Raman imaging utilizes inelastic light scattering to produce high-resolution chemical maps without the need for dyes or staining, and it has been deployed clinically for real-time cancer detection and distinguishing pathological from normal tissue^1^.

Depth-resolved Raman measurements have been used to characterize hydration gradients^2,3^ and lipid organization in the stratum corneum^4^. Raman imaging has also delineated biochemical differences between normal, dysplastic, and malignant skin lesions and monitored wound healing and inflammation through longitudinal changes in collagen, lipids, and other molecular constituents^5,6^. Despite the method’s capacity to probe physiological/pathological skin states with sub-micrometer resolution, spatial molecular profiling of skin microarchitectures remains underexplored.

Human skin comprises two principal layers—the epidermis and dermis—each harboring diverse cell populations with distinct molecular characteristics. Basal stem cells maintain tissue homeostasis in the interfollicular epidermis. Recent lineage tracing and single-cell studies in mice and humans have shown that epidermal stem cell populations are heterogeneous, exhibiting distinct gene expression profiles ^7–10^. Human skin is characterized by undulating epidermal–dermal junctions, forming rete ridges and inter-ridge regions corresponding to spatially segregated stem/progenitor niches^10–14^. To date, these niches have primarily been defined using antibody-based markers; truly label-free molecular signatures remain largely unknown.

Building on recent Raman-based identification of vascular disease biomarkers^15^ and single-cell Raman analysis^16^, in this study, we applied high-resolution Raman imaging combined with multivariate analysis to profile the molecular landscape of human skin. Examination of skin sections at single-cell resolution revealed spatially distinct molecular fingerprints delineating basal populations at the epidermal–dermal junction, defining stratified layers across the epidermis and dermis. These findings reveal the heterogeneity of human skin architecture and offer a label-free platform for assessing homeostasis, aging, and regenerative interventions.

## Results

Label-free Raman imaging was performed on unfixed, frozen tissue sections (Fig. 1A, B). Acquired spectra were analyzed using multivariate curve resolution–alternating least squares (MCR-ALS) and resolved into five distinct components: (1) an epidermal layer, (2) an epidermal basal layer, (3) a dermal layer, (4) nuclei, and (5) stratum corneum (Fig. 1C). Each component exhibited characteristic Raman peaks: proteins showed peaks at 1002, 1450, and 1660 cm^-1^; the nuclei component showed a strong peak at 787 cm^-117,18^. Lipid-rich regions such as the stratum corneum exhibited peaks at 1652 cm^-1^ (C=C stretching) and 1743 cm^-1^ (ester C=O stretching), indicative of lipid content^19.^ (Supporting Table 1). Heatmaps of each spectral component were generated based on the Raman intensity at characteristic peaks, providing a molecularly resolved visualization of human skin tissues (Fig. 1D).

**Fig. 1:**
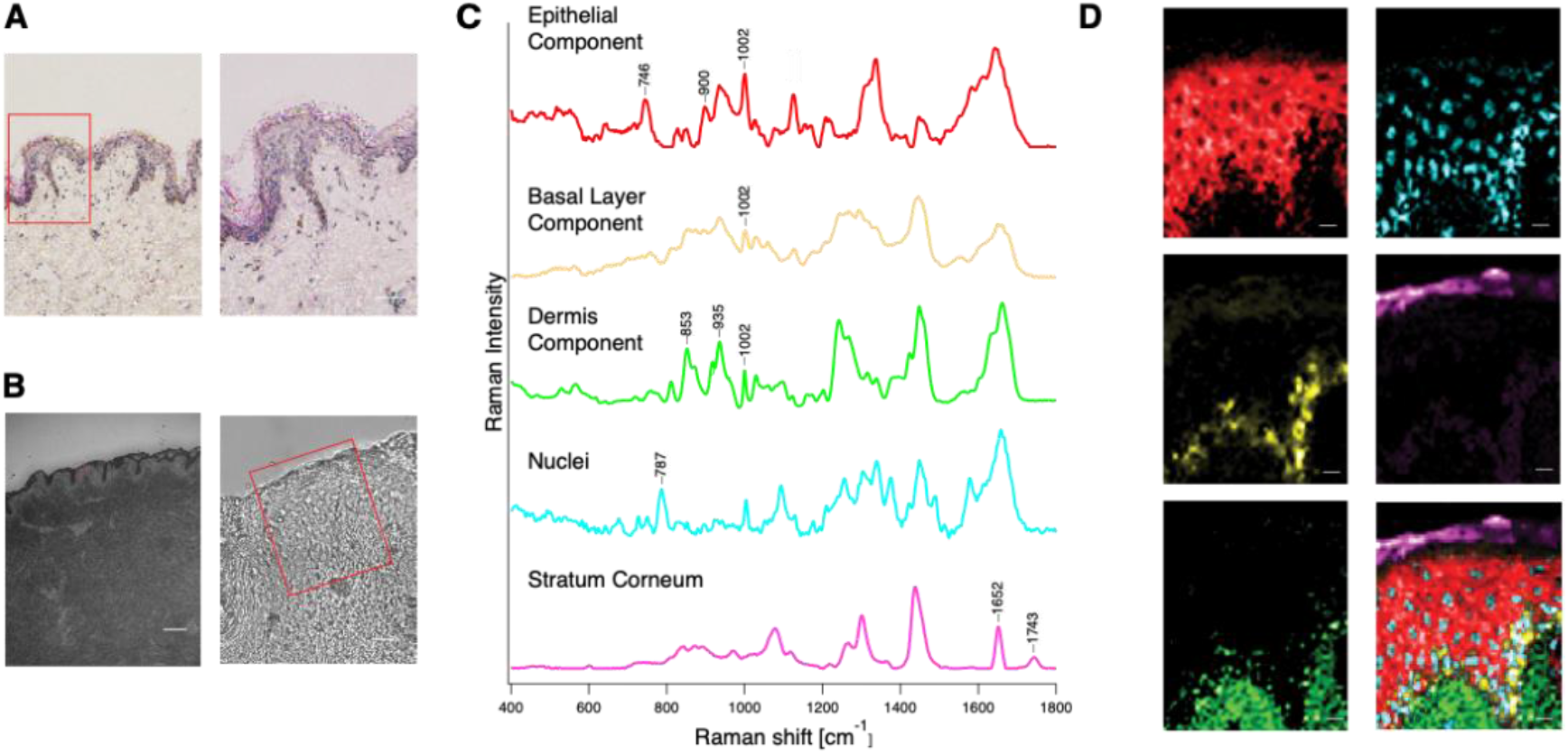
Label-free Raman imaging employed to visualize different cellular and molecular features in human skin. (A) Hematoxylin and eosin staining of young human skin. The left image shows a wider view (scale bar: 75 µm); the right image shows a magnified view (scale bar: 25 µm). An undulating epidermal–dermal structure is clearly visible in young skin. (B) Bright-field images acquired for Raman measurements. Red rectangles indicate regions of interest selected for Raman mapping. The left image shows a low-magnification field (scale bar: 100 µm); the right image shows a high-magnification field (scale bar: 20 µm). (C, D) Representative Raman spectra and the corresponding false-color Raman images of each component in human skin. Red: epithelial component; yellow: basal layer component; green: dermal component; blue: nuclei; magenta: stratum corneum lipids. Scale bars: 10 µm.

The nuclei spectra were further analyzed due to their localization in the basal region, where epidermal stem cells typically reside. Regions corresponding to the top and bottom of the rete ridge-like structures were manually segmented based on morphological features in the Raman intensity maps (Fig. 2 & Fig. 3). The spectra from these regions were employed in a principal component analysis (PCA) to identify the molecular features distinguishing the regions (Fig. 2A–D). The PC1–PC4 score plot revealed distinct clustering between top and bottom regions (Fig. 2B). The top region of the PC4 loading plot showed a higher score than the bottom; loading positive areas reflected the features of the top cells, with negative loadings corresponding to the bottom cells (Fig. 2C). The PC4 loading patterns revealed that the top region had peaks at 787 cm^-1^ for nucleic acids. The bottom region exhibited negative loadings at 1444 cm^-1^ for CH_2_/CH_3_ deformation and 1649 cm^-1^ for amide I, reflecting protein secondary structures (Fig. 2D). Positive loading areas reflected the features of the top cells; negative loadings corresponded to the bottom cells, suggesting that signals at the top of the undulations are characterized by protein-related features, while those at the bottom exhibit nucleic-associated spectral signatures. This indicates functional heterogeneity between the top and bottom regions of the undulating structure, potentially reflecting differences in epidermal stem cell characteristics and microenvironmental specialization.

**Fig. 2:**
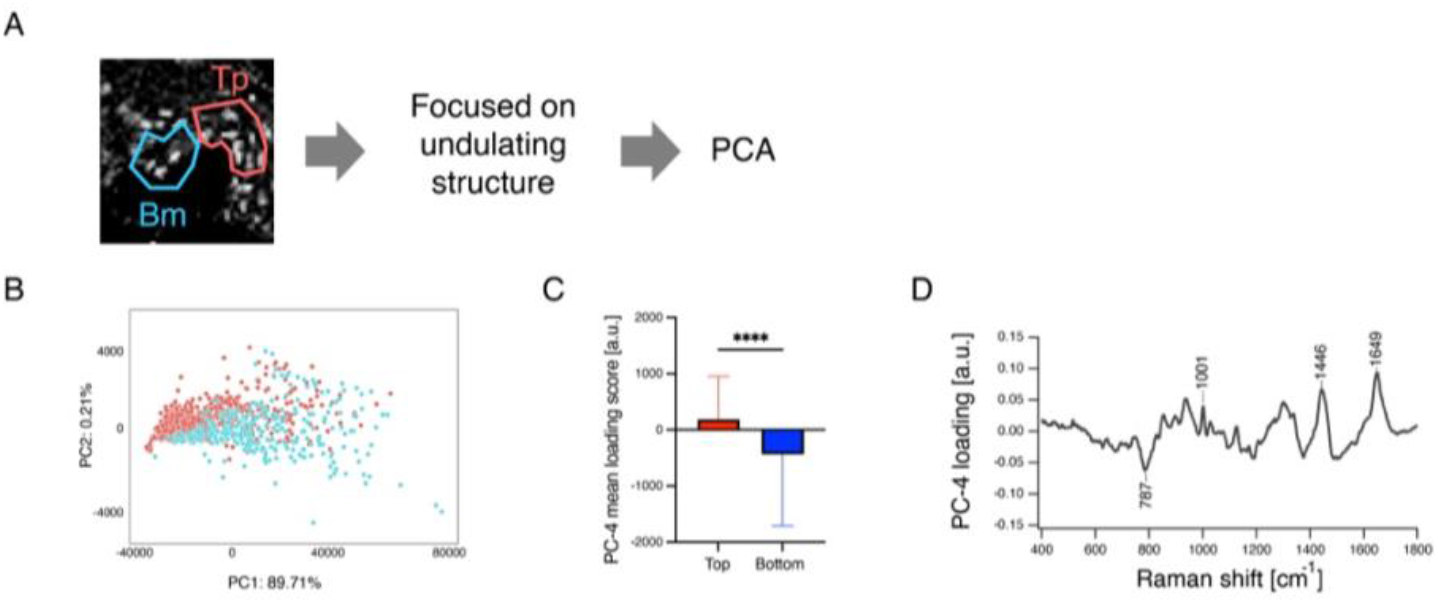
Determined molecular features for the undulating structures of human epidermis by PCA. (A) Raman spectra from the inter-ridge (top) and rete ridge (bottom) regions of epithelial cells were extracted from reconstructed Raman maps and subjected to principal component analysis (PCA). (B) Scatter plot of PC1 vs. PC4. Clustering of cells from the top (red) and bottom (blue) regions is clearly visible, with distinct separation along PC1 and PC4. (C) Mean loading scores of PC4 for top (red) and bottom (blue) cells. A Mann-Whitney U test revealed a significant difference between top and bottom cells (****P < 0.0001). (D) Loading plot of PC4. Peaks corresponding to nucleic acids are dominant in the negative direction; protein-related peaks are prominent in the positive direction.

**Fig. 3:**
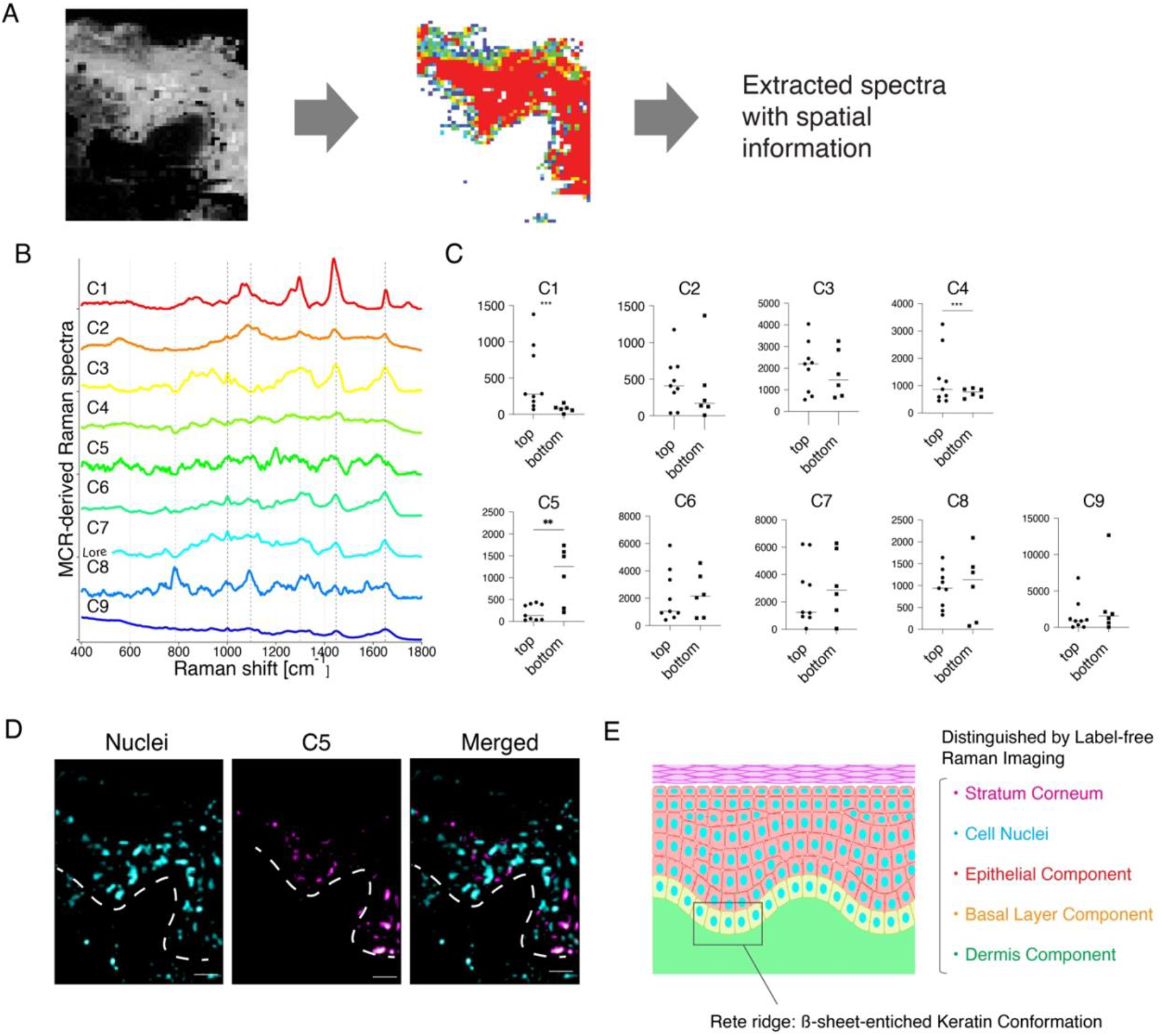
Label-free Raman imaging of molecular features within the undulating structure of human epidermis by MCR. (A) Further decomposition of the epithelial area was performed using multivariate curve resolution (MCR). Black-and-white images were used for masking; individual spectra were extracted from color-coded spatial domains. (B) MCR-derived Raman spectra focusing on the undulating skin structure, comparing the distributions between the top and bottom regions. Each component exhibits distinct Raman peaks: C1 = lipid component, C2–C9 = protein-containing components. (C) Statistical comparison of MCR-derived Raman intensities between top and bottom regions. Normality was assessed by the Shapiro–Wilk test. For normally distributed data, unpaired t-tests were applied; F-tests were used to compare variances. C1 and C4 showed significant variance differences (*** P < 0.001). The C5 component exhibited significantly higher intensity in the bottom region compared with the top (**P < 0.01). C1 and C4 were more abundant in the top region; C5 was enriched in the bottom region. (D) Raman image of the C5 component, which is specifically localized to the bottom region and not to the top. (E) Schematic summary of Raman imaging results in human skin.

Multivariate curve resolution (MCR) was performed to identify molecular biomarkers and distinguish each structure from the areas (Fig. 3A). The spectra were extracted from the epidermal region, followed by MCR. The spectral data were decomposed into nine spectra to explore region specificity (Fig. 3B). MCR derived nine components (C1–C9). Components C1–C4 demonstrated elevated expression in the top region, with statistically significant differences observed between C1 and C4. Components C5–C9 were enriched in the bottom region, and C5 exhibited a significant difference. The C1 spectrum showed lipids at 1297, 1437, and 1652 cm^-1^; C2, C3, C4, C5, C6, C7, and C9 showed proteins at 1002, 1445, and 1650 cm^-1^, and C8 showed nucleic acids at 782 and 1092 cm^-1^ (Supporting Table 2).Notably, C5 exhibited distinct peaks at 1200 and ∼1350 cm^-1^, characteristic of keratin from the backbone and side-chain vibration of keratin molecules^20^.

The comparison revealed a statistically significant increase in the C5 signal in the bottom region compared to the top (Fig. 3C), indicating that this component is preferentially expressed in the bottom part of the undulating structure. Raman-based spatial reconstruction of the C5 and nuclei components confirmed preferential localization of C5 to the bottom of the undulating structure (Fig. 3D), suggesting that it may serve as a potential molecular marker for this region.

In C5, the amide III band exhibited a distinct shift toward lower wavenumbers, with a main peak around 1237 cm^-1^ and a broad shoulder near 1240 cm^-1^, compared to spectra representing the top areas. This downshift is indicative of reduced α-helical content and increased β-sheet or extended conformational content. Previous Raman spectroscopic studies of keratin and other fibrous proteins have reported similar low wavenumber shifts accompanying secondary structure transitions, where α-helix-dominated spectra (∼1260 cm^-1^) evolve into β-sheet-rich bands near 1240 cm^-118^.

The observed lower shift in the amide III band and broadening of the C5 component can be interpreted as spectral signatures reflecting β-sheet-enriched or flexible keratin structures present in the basal region of epidermal undulations (Fig. 3E). These structural features likely represent an adaptive molecular organization contributing to plasticity of the basal epidermis and its capacity for renewal^18–20^.

## Discussion

Label-free confocal Raman imaging enabled molecular resolution of the stratum corneum, all viable epidermal layers, the collagen-rich dermis, and individual keratinocyte nuclei, providing a comprehensive molecular map of human skin microarchitecture (Fig. 1C, D).Raman microspectroscopy, combined with multivariate analysis, identified protein-and polysaccharide-derived spectral features specifically associated with the undulating architecture of the epidermal–dermal junction. These biochemical markers coincided with regions enriched in epidermal stem cells, suggesting their potential utility as label-free indicators of the functionally specialized undulating structures within the epidermis.

Raman imaging, which enables the molecular-level examination of conventional *in vitro* skin models, is widely used in pharmacological testing and regenerative medicine. The spectral markers identified in this work provide a blueprint for engineering skin constructs that faithfully reproduce the epidermal–dermal undulations essential for stem cell localization and function. While Raman microscopy has been employed to monitor tumor organoid growth *in vivo*^21^, no study to date has applied the technique for tracking longitudinal molecular changes in skin models while preserving information on the undulating epidermal–dermal interface.

This analysis focused on the undulating epidermal–dermal interface, contrasting spectra obtained from the inter-ridge (top) and rete ridge (bottom) regions (Fig. 2-3). Lasso regression with MCR was applied to pinpoint the spectral features that distinguish these microdomains. The analysis identified component C5, which was selectively enriched in the bottom regions (Fig. 3D). The C5 profile exhibited the structural signature of keratin, suggesting its potential as a label-free biomarker for the basal segment of the undulating structure.

Raman spectroscopy can resolve differences in molecular secondary structure rather than simply detect molecular abundance. The method revealed that areas at the basal side of the epidermal undulations that exhibited an amide III band centered around 1237 cm^-1^, indicating a higher proportion of β-sheet or extended keratin conformations^18^ and suggesting that keratin filaments in the basal region adopt a less tightly coiled configuration than the canonical α-helical form. Such β-sheet-enriched keratin structures may represent a localized structural adaptation within the basal epidermis, contributing to the flexibility and resilience necessary for tissue maintenance and renewal.

As Raman-detectable signatures can be captured without tissue processing, they represent an objective, non-invasive gauge of how faithfully *in vitro* skin models reproduce the native epidermal–dermal topography. Importantly, the non-destructive nature of Raman imaging allows for revisiting the same construct, enabling the charting of the appearance, maturation, and long-term stability of rete ridge-associated markers during culture. Such longitudinal feedback could streamline the optimization of engineered skin equivalents, facilitate high-content drug-screening assays that require an intact barrier architecture, and accelerate the translation of tissue-engineered grafts by delineating the biomechanical and stem cell niches of native skin.

The ability to evaluate and refine skin equivalents based on molecular signatures associated with epidermal stem cell niches is promising in regenerative medicine.Integrating these identified Raman-derived markers into the design/quality control pipelines of skin substitutes or cell sheets could enable grafts with superior functional integration and long-term stability. These findings therefore advance the development of label-free, quantitative criteria for assessing stem cell-rich architectures in skin tissue engineering.

## Methods

### Human skin samples and ethics

Frozen full-thickness normal human abdominal skin samples from donors aged 30s to 40s were purchased from CTI-Biotech (Lyon, France). OCT compound (Tissue-Tek, Sakura Finetek) was used to snap-freeze human skin. Informed consent was obtained from anonymous donor patients for the collection of these samples. The consent and collection procedures were compliant with the European standards and applicable local ethical guidelines.

### Hematoxylin and eosin staining

For the hematoxylin and eosin staining, 10-mm skin sections were air-dried, fixed in 4% paraformaldehyde at room temperature for 10 min, and washed in PBS. The sections were stained with hematoxylin (Wako) for 20 min and eosin Y (Wako) for 5 min. After dehydration, the sections were mounted using Entellan’s new mounting solution (Merck Millipore). Images were captured using the EVOS M5000 Imaging System (Thermo Fisher Scientific, Waltham, MA, USA).

### Raman measurements

Raman measurements were obtained using confocal microscopy (Xplora Plus, Horiba, Kyoto, Japan) equipped with a 532-nm laser with 30 mW of power. Cross sections of human skin on glass slides were rinsed with PBS, and data acquisition was performed using a 60× apochromat water dipping objective (N.A. 1.1, Olympus). High-resolution Raman imaging was performed in areas measuring 100 × 200 µm with 2 × 2 µm pixel resolution and 0.5 s acquisition per spectrum.

### Data analysis

#### Denoised Raman spectra

The obtained Raman spectra were processed using IGOR Pro software (WaveMetrics, Inc., Lake Oswego, OR). Indene spectra were employed for the calibration of the standard wavelength. The halogen lamp spectra were utilized to calibrate both the detector and the intensity, which were adjusted using the white light spectrum. Cosmic rays were removed if present. The wavenumber calibration was conducted by fitting the indene Raman spectra peak positions with fourth-order polynomial functions. For noise reduction, singular value decomposition (SVD) was employed to reconstruct the spectra using 20 SVD components; this was carried out prior to the MCR-ALS analysis.

#### Multivariate curve resolution

MCR-ALS was applied to extract molecular-based spectra^22^. Following the denoising procedure, human skin-derived Raman spectra were preprocessed using MCR-ALS to eliminate the autofluorescence background^23^. MCR-ALS was optimized for biological tissues using Raman spectroscopy measurements^15^. SVD initialization was used to identify MCR components for the murine analysis. The MCR results from the murine study were utilized as initial spectra for the human analysis. The MCR-ALS calculations were performed using the SciPy library in Python.

#### Spectra extraction

A targeted extraction approach based on intensity information from MCR-ALS component analysis was implemented to analyze specific regions of interest within the tissue sections. The intensity distribution of each MCR-derived component was visualized as a heatmap across the entire tissue section. Reference areas known to lack the component of interest (such as areas outside the tissue boundary) were analyzed to establish appropriate threshold values distinguishing signal from background noise.

Binary masks were generated for each component by applying these thresholds, creating a binary representation where positive signals were retained and noise was excluded. These masks enabled the extraction of spectra exclusively from regions where the component of interest was present at significant levels. For the analysis of the undulating structures, separate masks were created for the top and bottom regions based on their spatial location within the epithelial layer, utilizing the intensity of nuclei-derived components. For epithelial areas, protein-derived components were used for extraction. The extracted spectral datasets maintained their spatial information, allowing correlation between molecular signatures and histological features. The extracted spectra then underwent further statistical analysis.

#### Principal component analysis

Raman spectral datasets corresponding to MCR-derived components were extracted and subjected to PCA using the nonlinear iterative partial least squares (NIPALS) algorithm. PCA, widely used in chemometrics, facilitates the identification and interpretation of the spectral variation within a Raman dataset^24^. PCA represents spectral information as vectors termed principal components (PCs). The results are typically visualized using a scores plot, where one principal component is plotted against another and each spectrum is represented by an individual point. The corresponding PC loadings plot highlights the peaks that significantly influence the score values, enabling interpretation of molecular differences between groups^25^. Data analysis was performed using the Scikit-learn package in Python.

### Statistical analysis

Data are presented as mean ± SD. Statistical analyses were carried out using Prism 10 software (GraphPad, La Jolla, CA, USA). The Shapiro–Wilk test was used to assess the normality of the data. For normally distributed data, statistical significance was evaluated using t-tests (paired or unpaired) for two-group comparisons and one-way or two-way analysis of variance for comparisons between three or more groups, followed by the Bonferroni correction for multiple comparisons. Nonparametric tests such as the Mann– Whitney U or Kruskal–Wallis tests were applied when the data did not meet the normality assumption. A P-value of 0.05 or less was considered statistically significant.

## Data availability

Available upon request.

## Code availability

Available upon request.

## Acknowledgements

We thank the International Core-facility of Advanced Life Science at Kumamoto University and the Research Promotion Unit at Kyushu University for their invaluable support. We also thank T. Keida for her technical assistance with H&E staining. This work was supported by the Program for Technological Innovation of Regenerative Medicine, AMED (23bm0704067) (to AS and KS), Kose Cosmetology Research Foundation from Kose Cosmetic Co.(to KS), the project of JPNP14004, commissioned by the New Energy and Industrial Technology Development Organization (NEDO) from (to MA), the IKEDARIKA Grant from Ikedarika Scientific Co., Ltd., and Leave a Nest Co., Ltd. (to MI). This work was supported in part by the MEXT Cooperative Research Project Program, the Medical Research Center Initiative for High-Depth Omics, and CURE: JPMXP1323015486 for MIB, Kyushu University.

## Author contributions

KS, MA, AS, and HT: experimental design. MI and AS: skin preparation and discussion. KS and MA: Raman measurements and analysis. KS, MI, and AS: skin parts, mainly discussed. All authors are discussed.

## Competing interests

None.

## Notes

### Competing Interest Statement

The authors have declared no competing interest.

